# Adapting to visuomotor rotations in stepped increments increases implicit motor learning

**DOI:** 10.1101/2022.07.04.498746

**Authors:** Shanaathanan Modchalingam, Marco Ciccone, Sebastian D’Amario, Bernard Marius ’t Hart, Denise Y.P. Henriques

## Abstract

Human motor adaptation relies on both explicit conscious strategies and implicit unconscious updating of internal models to correct motor errors. Implicit adaptation is powerful, requiring less preparation time before executing adapted movements, but recent work suggests it is limited to some absolute magnitude regardless of the size of a visuomotor perturbation when the perturbation is introduced abruptly. It is commonly assumed that gradually introducing a perturbation should lead to improved implicit learning beyond this limit, but outcomes are conflicting. We tested whether introducing a perturbation in two distinct gradual methods can overcome the apparent limit and explain past conflicting findings. We found that gradually introducing a perturbation in a stepped manner, where participants were given time to adapt to each partial step before being introduced to a larger partial step, led to ∼80% higher implicit aftereffects of learning, but introducing it in a ramped manner, where participants adapted larger rotations on each subsequent reach, did not. Our results clearly show that gradual introduction of a perturbation can lead to substantially larger implicit adaptation, as well as identify the type of introduction that is necessary to do so.

## INTRODUCTION

The human motor system can quickly adapt to errors in our movements to improve future performance. Motor adaptation involves multiple learning processes including explicit conscious strategies that change our movement intent and implicit updates to internal models controlling the execution of our intended movements [1–6]. Although the interaction between the two processes is crucial to motor learning, studying them in isolation lets us glean how each may uniquely contribute to our ever-changing motor repertoire. In this study, we focus on implicit motor learning and its apparent limits. We show that our current understanding of implicit motor learning cannot fully explain its occasionally varying asymptotic limits in visuomotor rotation paradigms and provide a method of increasing implicit contributions to motor learning.

In many visuomotor adaptation paradigms, adapting reaching movements to consistent perturbations eventually leads to implicit learning reaching a steady-state below full compensation of the perturbation [7–10]. When people adapt to relatively large perturbations, we repeatedly observe an upper bound to asymptotic implicit learning that is independent of the perturbation size [7,9]. This effect is observed when implicit learning is measured by subtracting planned strategy use [7], by restricting strategy use after adaptation [11,12], or when people implicitly adapt to invariant motor errors [9,13]. Thus, this cap in implicit adaptation appears robust.

The factors governing the upper limits of implicit learning are an area of active study [9,14,15]. While other errors may influence implicit learning [14,16,17], it is considered largely driven by sensory prediction error (SPE) [16,18,19] – that is, the difference between the predicted sensory consequences of a planned movement and the perceived sensory consequences upon movement execution. When adapting to large perturbations that are introduced abruptly, participants may become aware of a perturbation and explicit re-aiming strategies may be formed to partially counter the perturbation. Since the magnitude of explicit strategies is flexible and scales to the size of the perturbation, larger perturbations can bring about additionally reduced implicit motor adaptation [7,11,12], resulting in apparent limits to implicit learning.

Conversely, it follows that limiting explicit strategy use should increase implicit learning. By introducing the perturbation gradually, we can limit any noticeable errors for participants to construct explicit strategies. This should increase implicit learning. Some studies found this [20–23], but in other studies measures of implicit learning are similar regardless of whether the perturbation was introduced abruptly or gradually [24–26]. While some of the discrepancies could be due to failure to exclude strategies in measures of implicit adaptation, the primary and untested difference could be how gradual perturbations are introduced.

Perturbations are typically gradually introduced to participants using one of two ways: in a ramped manner where the magnitude of the perturbation is increased a small amount every trial, or in distinct steps where participants experience the same perturbation for multiple trials before the perturbation is further increased. For example, in the seminal study by Kagerer et al. [20], participants adapted to perturbations in 10° incremental steps, each step consisting of 60 reaches. In other studies, participants adapted to perturbations in a ramped manner, where perturbations increased by small amounts on each subsequent reach [21,27]. Here, we directly compare the effect of these two types of gradual introduction on implicit learning.

We also attempted to isolate implicit learning during classical visuomotor adaptation, as well as measure explicit strategy use in a comparable way, by using a strategy exclusion method known as the process dissociation procedure [5]. In this way, we tested whether introducing the perturbation more gradually (whether ramped or stepped), and thus purportedly reducing explicit strategy by providing smaller, less salient errors does elicit higher asymptotic implicit motor learning. We show that it can, but only when perturbation is introduced in a stepped manner which suggests that not all gradual introductions are equal.

## METHODS

### Participants

105 participants participated in the study. All participants self-reported to be right-handed and had normal or corrected-to-normal vision. Participation was voluntary and all participants gave informed, written consent prior to data collection. The procedures used in this study were approved by the York Human Participants Review Sub-committee.

### Setup/Apparatus

Participants sat on a height-adjustable chair facing an experimental apparatus containing a downward-facing LCD screen (30Hz, 20″, 1680 × 1050, Dell E2009Wt) and mirror to display visual feedback, and a tablet and pen (30Hz, 1920 × 1080, Wacom Intuos Pro PTH860) to record the hand position (Fig 1A). They individually adjusted the chair’s height and distance from the apparatus until the display was fully visible and they could comfortably make reaching movements across the tablet. Participants held the pen in a precision grip and maintained contact with the tablet throughout the experiment. A thick cloth draped over their right shoulder (not shown in Fig 1) occluded vision of their arm movements. All visual feedback was presented through the downward-facing screen reflected by the mirror. The position of the mirror projected the visual feedback to appear as if on the same horizontal plane as the participants’ hand movements.

**Fig 1:**
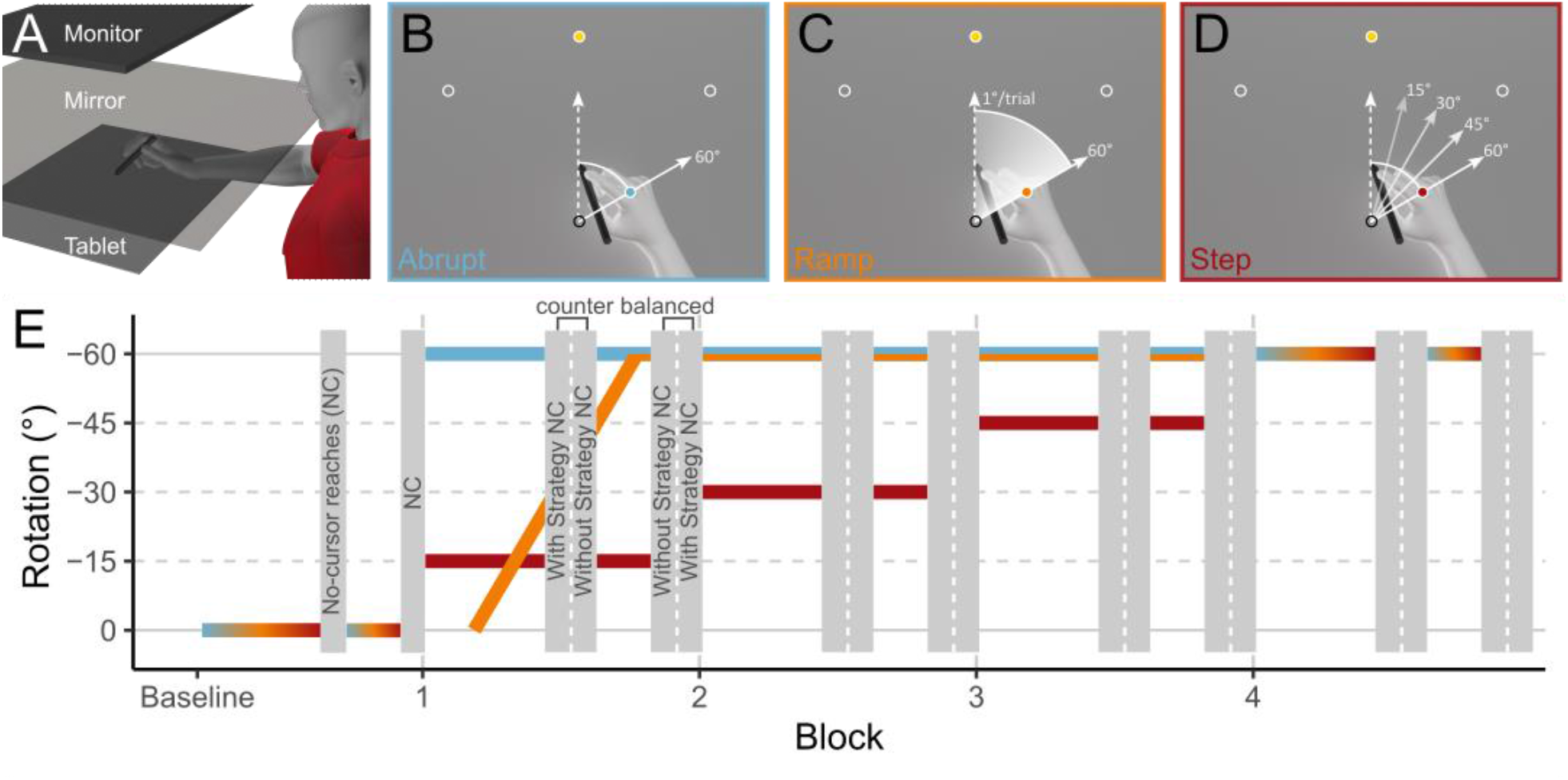
A: Experimental apparatus for task presentation (monitor and mirror) and data collection (tablet) B-D: Methods of perturbation introduction for the three groups of participants. E: The schedule of tasks presented to participants over the course of the experiment.

### Visuomotor rotation tasks

Participants made outward reaches towards 1 cm-diameter circular targets by smoothly gliding the pen held in their hand along the touchpad. The targets were located 10 cm away from the starting position at one of 3 possible locations: 45, 90, or 135 degrees in polar coordinates (Fig 1: B-D). Participants were instructed to make quick, straight reaches towards the target but were not penalized for failing to do so. In tasks where participants received visual feedback of movements, called “Reach-with-cursor” tasks, a 1 cm-diameter circle represented their hand position. To complete a Reach-with-cursor task, the center of the hand cursor needed to be within 0.5 cm of the target’s center, while remaining still on the touchpad for 500ms. In other tasks, called “No-cursor” tasks, visual feedback of the hand was not provided during or after the movement. To complete No-cursor tasks, participants needed to indicate the completion of their movement by holding their hand still for 500ms following a reach. For these tasks, they were not required to acquire the target. After completion of either task, participants were instructed to move back to the starting position to begin the next trial.

### Experiment protocol

The participants were randomly assigned to 3 possible groups: “Abrupt”, “Ramp” and “Step”. Each group experienced a different method of perturbation introduction. Figure 1: B-D shows the perturbation experienced in Reach-with-cursor tasks by each group throughout the experiment. In trials where the perturbation was 0°, the position of the hand-cursor was aligned with that of the participant’s real hand. When a non-zero perturbation was applied, any movement along the touchpad was rotated by the magnitude of the perturbation.

All three experiments consisted of 5 blocks, divided into two phases: the “Baseline” phase and the “Rotated” phase. A block began with the Reach-with-cursor task (45 trials), followed by the No-cursor task (9 trials in the Baseline phase and 18 trials in the Training phase), followed then by the Reach-with-cursor task (21 trials), and finally by the No-cursor task (9 trials in the Baseline phase and 18 trials in the Training phase). When transitioning between tasks, the words “Reach to Target” and “Reach with No Cursor” were displayed to participants to indicate the next trial belonging to the Reach-with-cursor task and No-cursor task respectively. The Baseline phase consisted of one block and the rotated phase consisted of 4 blocks.

Participants took a short break before starting the Rotated phase. During the break, participants were told there may be changes in the behaviour of the hand cursor for the rest of the experiment. Participants were told to mentally note any strategy they may employ to counter the behaviour of the hand-cursor, as they may be asked to use or not use that strategy during No-cursor tasks. Detailed instructions can be found at https://osf.io/a7k6s/.

To isolate implicit aftereffects of learning, and to assess explicit strategy use following adaptation, we used a process dissociation procedure (PDP). During the Rotated phase of the experiment, in addition to the “Reach with No Cursor” instructions, the words “WITH strategy” or “WITHOUT strategy” cued participants to employ, or not employ any cognitive strategy they may have used for the next 9 No-cursor trials. After completing the 9 trials, participants received the alternate instruction for the next 9 No-cursor trials. The order of the With and Without Strategy instructions was counterbalanced to avoid order effects. We used deviations from baseline performance in Without-strategy-no-cursor tasks as our measure of implicit aftereffects of learning. As our measure of explicit strategy use, we used any added deviation evoked by our instruction to use a remembered strategy.

### Analysis

To measure performance during our reaching tasks, we calculated the angular deviation of a participant’s hand path from a straight-line movement to the target at the point of maximum velocity – which we term “hand deviation”. When reaching with a rotated cursor in the Rotated phase of the experiment, participants had to deviate their hand paths counterclockwise by the full amount of the visuomotor perturbation to make the cursor move directly to the target. We corrected for individual biases in reach behaviour by subtracting the mean hand deviations for each target in the Baseline phase of the experiment from hand deviations to the corresponding targets in the rotated phase. All subsequent calculations and data analysis were conducted on baseline-corrected data. To determine if participants adapted to the visuomotor rotation presented to them, we analyzed performance during key trial sets during Reach-with-cursor tasks in the Rotated phase of the experiment. The first trial set included the first 3 trials during learning, representing one reach to each of the three possible targets with the perturbation applied. Only the participants in the Abrupt group experienced the full 60° rotation during this trial set. Additional trial sets of interest included the final 6 Reach-with-cursor tasks in Block 1 for the Abrupt and Ramp groups and the final 6 Reach-with-cursor tasks in Block 4 for all groups.

To confirm that participants in all groups adapted to the 60° rotation, we conducted a 2×3 mixed analysis of variance (ANOVA) with the group which participants belonged to as a between-subject factor and the Initial and Final block trial sets of interest (See Fig2A: grey-shaded areas) as a within-subject factor.

To test whether the method of perturbation introduction affected implicit motor adaptation, we compared the aftereffects of learning as measured in Without Strategy No-cursor trials. We made two sets of comparisons. First, we compared hand deviations during these No-cursor tasks in the initial blocks in which each group first became exposed to the full 60° rotation: Block 4 for the Step group, and Block 1 for the Ramp and Abrupt groups. Additionally, to account for time spent adapting to a visuomotor perturbation, we compared Hand deviations during Without Strategy No-cursor tasks in Block 4 for all groups. For both comparisons, we first calculated mean hand deviations during each block for each participant. We then performed a one-way ANOVA to determine the effects of the method of perturbation introduction on hand deviations during these Without Strategy No-cursor tasks. For all comparisons, we computed Inclusion Bayes factors between models that include or do not include relevant effects [28]. To analyze strategy use following adaptation, we calculated the mean explicit strategy used during each block by each participant. We did this by subtracting mean hand deviations within a block during Without Strategy No-cursor tasks from mean hand deviations within a block during With Strategy No-cursor tasks. We repeated the analyses specified above for the explicit strategies participants employed.

We also explored if the method of perturbation introduction affected the temporal dynamics of implicit aftereffects of learning and explicit strategy use – both measured via the PDP. We expected implicit aftereffects to decay over time. We calculated mean rates of change in hand deviations during each block of No-cursor tasks for participants in the rotated phase of the experiment. For each participant, we fitted a simple linear model predicting hand deviation as a function of the trial number within a No-cursor block. To determine the effects of the method of perturbation introduction and the block number on decay rates of implicit and explicit motor adaptation processes, we conducted separate two-way 3 (method of perturbation introduction) x 4 (block) mixed ANOVAs for implicit and explicit effects of learning, with the method of perturbation introduction as a between-subject factor and block as a within-subject factor.

## RESULTS

Participants in all groups adapted to the presented perturbations over the 4 blocks of the rotated phase of the experiment (main effect of trial set: F_(1,102)_ = 1681.30, p < .001, η^2^_G_ = .87, BF_incl_ > 10^6^) (Fig 2: Init vs trial set “4”). The method of perturbation introduction further affected the extent of learning in the Rotated phase (interaction between group and trial set: F_(2, 102)_ = 15.48, p < .001, η^2^_G_ = .11, BF_incl_ > 10^6^). This is likely driven by differences in the initial trials of the rotated phase of the experiment, as by the end of the final block, participants in all groups were able to adapt to ∼86% of the 60° visuomotor rotation (Step group mean: 51.81°, 86.35%; Ramp group mean: 50.34°, 83.90%; Abrupt group mean: 53.02°, 88.37%). We find moderate evidence that all three groups adapted to a similar level by block 4 of the experiment (BF_incl_ = 0.30).

**Fig 2:**
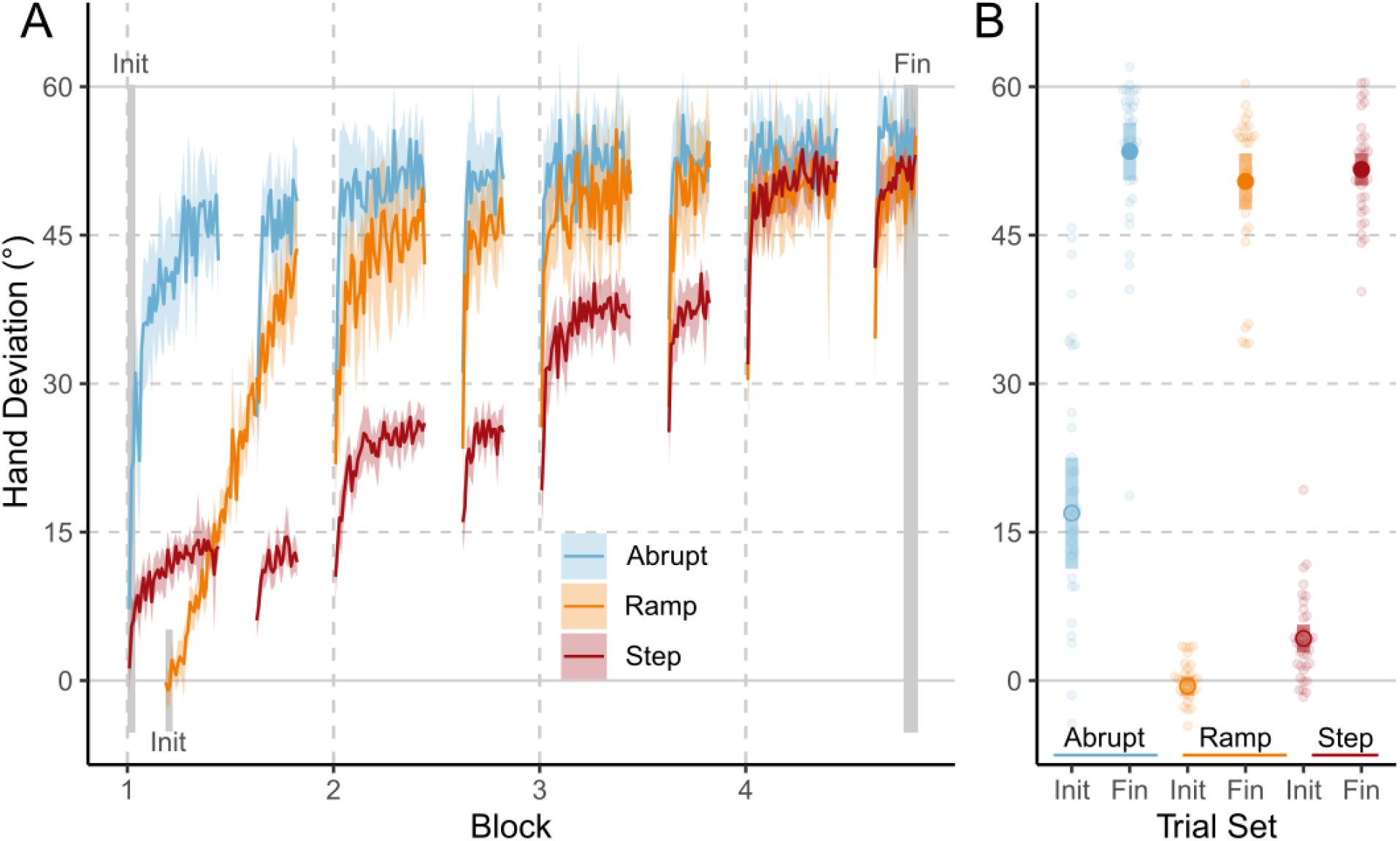
A: Mean hand deviations from straight-to-target reaches during Reach-with-cursor trials. B: Hand deviations in trial sets of interest. Coloured regions represent 95% CI.

To measure implicit aftereffects of adaptation, we asked participants to exclude any explicit strategies they used to compensate for the perturbation during training. During the first block of reaching with a 60° visuomotor rotation (Block 1 for the Abrupt and Ramp groups, Block 4 for the Step group), aftereffects of learning were affected by the method of perturbation introduction (Fig 3A: main effect of group: F_(2, 102)_ = 44.20, p < 0.001, η^2^ = .46, BF_incl_ > 10^6^). Implicit learning was higher for participants in the Step group (mean = 21.96°, sd = 8.07°) than those in the Ramp group (mean = 10.74°, sd = 3.71°, Tukey’s HSD: p < .001, BF_10_ > 10^6^) and the Abrupt group (mean = 10.76°, sd = 4.50°, Tukey’s HSD: p < .001, BF_10_ > 10^6^). We found moderate evidence that participants in the Ramp and Abrupt groups have similar amounts of implicit learning (BF_10_ = 0.249) after one block of adaptation.

**Fig 3:**
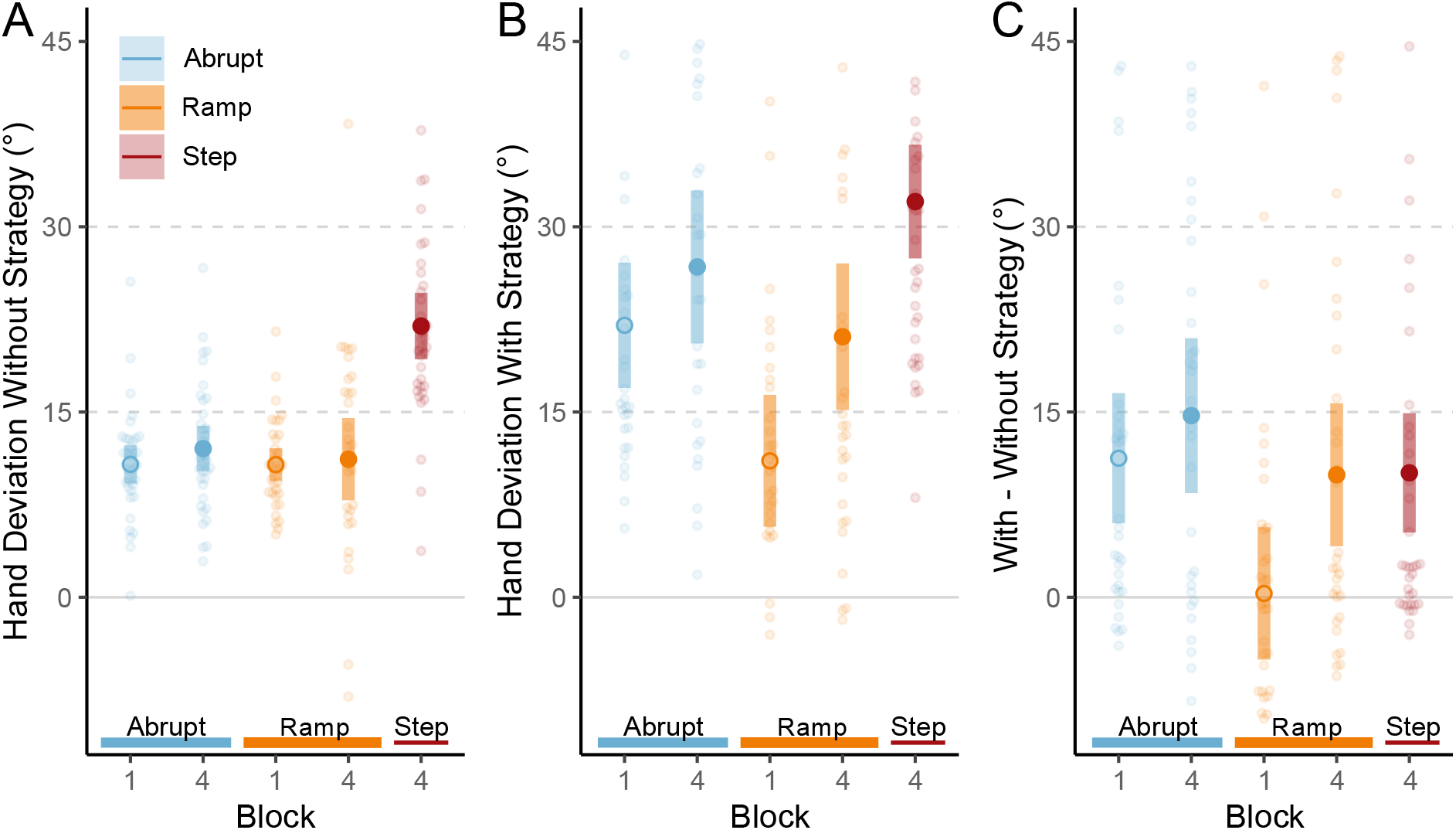
Mean hand deviations from straight-to-target reaches during No-cursor tasks without (A) and with (B) strategy use. C: Mean explicit strategies (calculated by subtracting Without-strategy hand deviations from With-strategy hand deviations). Coloured regions represent 95% CI.

To account for time spent adapting to a visuomotor perturbation, we also repeated our analyses on the final block of each condition. The method of perturbation introduction affected the implicit aftereffects in the final block of the experiment (Fig 3A: main effect of group: F_(2, 102)_ = 21.43, p < 0.001, η^2^ = .30, BF_incl_ = 7.33×10^5^). Post hoc tests revealed that after 4 blocks of adaptation, participants in the Step group exhibited ∼80% higher implicit learning (mean = 21.96°, sd = 8.07°) than those in the Ramp group (mean = 11.18°, sd = 9.38°, Tukey’s HSD: p < .001, BF_10_ = 6.57×10^3^) and the Abrupt group (mean = 12.04°, sd = 5.34°, Tukey’s HSD: p < .001, BF_10_ = 2.20×10^5^). We find moderate evidence that participants in the Ramp and Abrupt groups have similar amounts of implicit learning after being exposed to the perturbation for 4 blocks (BF_10_ = 0.27).

When initially experiencing a 60° perturbation – after 1 block in the Ramp and Abrupt groups, and after 4 blocks in the Step group – explicit strategy use was affected by the method of perturbation introduction (Fig 3C: main effect of group: F_(2,102)_ = 5.46, p = 0.005, η^2^ = 0.10, BF_incl_ = 6.80). During these initial blocks of experiencing a 60° perturbation, there was moderate evidence that participants in the Step and Abrupt groups could evoke strategies to account for the same proportion of a 60° perturbation (BF_10_ = 0.26). Furthermore, after 4 blocks of training, we found moderate evidence of similar explicit strategy use when cued in all groups (Fig 3B: BF_incl_ = 0.19)

Both implicit aftereffects of learning, measured via the exclusion of explicit strategies during No-cursor reaches, and explicit strategies may decay over time when repeatedly reaching without a cursor. To test whether these decay rates were affected by the method of perturbation introduction, we compared the rate of change in performance during 9-trial No-Cursor tasks. The method of perturbation introduction affected the rate of change of implicit aftereffects during 9-trial set of No-Cursor reaches (Fig 4A: main effect of group F_(2,102)_ = 7.36, p = 0.001, η^2^ = 0.05, BF_incl_ = 11.51) but the block number did not (F_(3,306)_ = 1.01, p = 0.387, BF_incl_ = 0.03). The rate of change of explicit strategies was not affected by either the method of perturbation of introduction (Fig 4C: no effect of group F_(2,102)_ = 0.95, p = 0.389, BF_incl_ = 0.06) or the block number (F_(3,306)_ = 1.08, p = 0.358, BF_incl_ = 0.03). When collapsed among blocks participants in the Step group had lower rates of decay over 9 No-cursor reaches (mean = -0.07°/trial, sd = 0.53°/trial) than those in the Ramp group (mean = -0.51°/trial, sd = 0.62°/trial, Tukey’s HSD: p = .003, BF_10_ = 17.37) and the Abrupt group (mean = -0.49°/trial, sd = 0.48°/trial, Tukey’s HSD: p = .005, BF_10_ = 31.83). Our exploratory analysis on the rates of change of implicit and explicit components of learning suggests a possible mechanism by which stepped introduction of perturbations led to increased implicit aftereffects, but a more thorough examination of decay rates over a larger number of No-cursor reaches is necessary.

**Fig 4:**
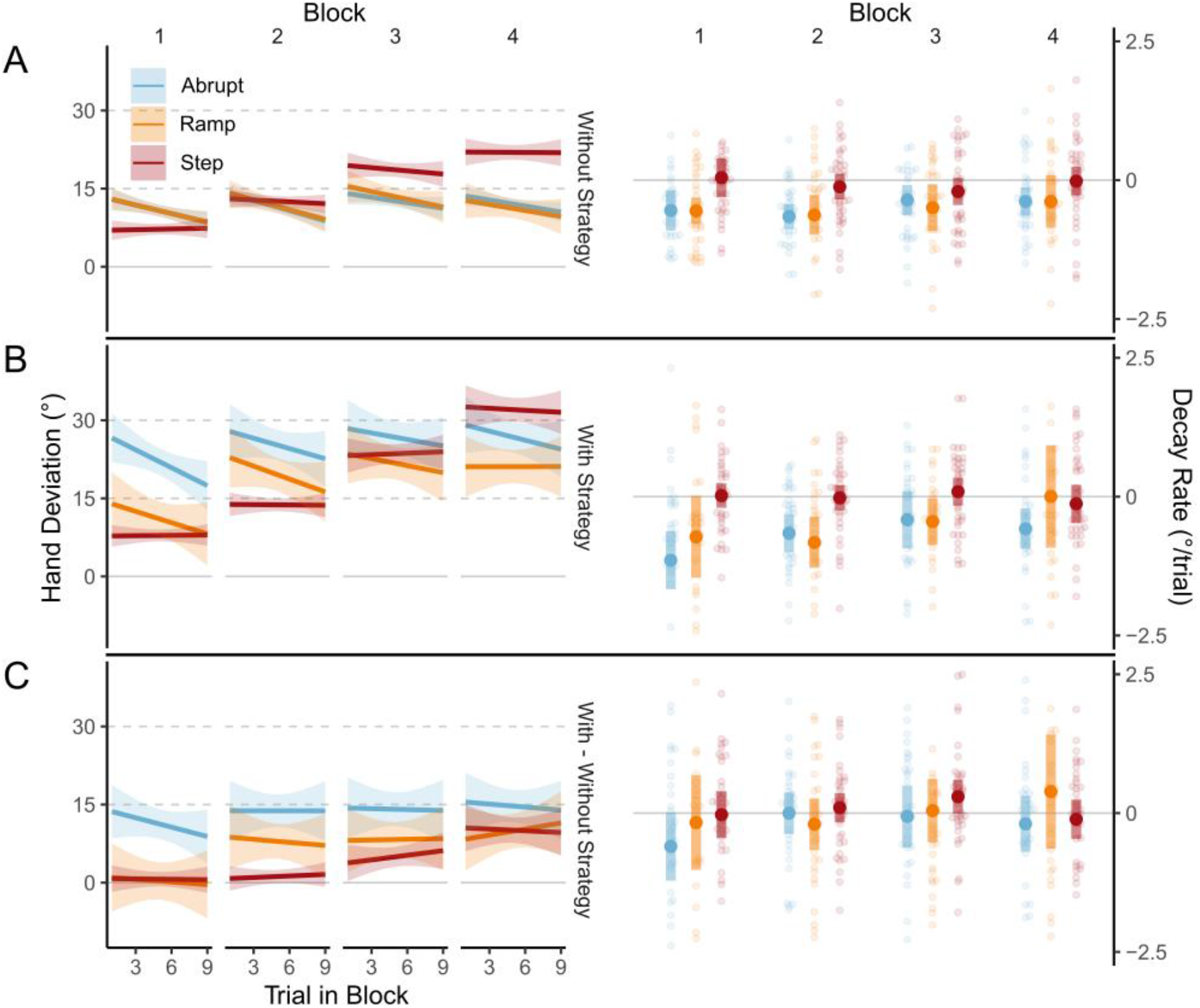
Left column: Mean hand deviations from straight-to-target reaches and calculated explicit strategies over 9 consecutive trials during No-cursor tasks without (A) and with strategy use (B), and calculated explicit strategies (C). Right column: Rates of change over 9 consecutive trials of right-column values during each block of the Rotated phase. Coloured regions represent 99% CI.

## DISCUSSION

The extent of asymptotic implicit adaptation in visuomotor rotation reaching paradigms is commonly held to have an upper limit independent of the size of the perturbation when a perturbation is abruptly introduced. It is also thought that this limit may be overcome by gradually introducing the perturbation. Here, we find that gradually introducing a perturbation in a stepped manner, but not in a ramped manner, led to ∼80% larger implicit aftereffects of learning, compared to abruptly introducing the perturbation.

We reproduced the often observed ∼15° limit to implicit motor adaptations when adapting to a large, abruptly introduced visuomotor rotation – measured with strategy exclusion trials. Whether this cap also applies to perturbations that are less salient, like those gradually introduced, is unsettled. Some previous studies that gradually introduced perturbations show possibly larger aftereffects [20,21,27].

Furthermore, recent studies have proposed that implicit adaptation processes both compete with and compensate for explicit strategies [14,29]. There is consequently strong theoretical backing for increased implicit learning in response to gradually introduced perturbations. However, there are many methods of introducing a perturbation gradually, and although they are generally considered under the same umbrella, it is not clear whether all have similar effects on implicit, or even explicit, learning. Against our expectations, when we directly compare these two types of gradual learning, we found increased implicit aftereffects of learning only when perturbations were introduced in a stepped, but not a ramped manner. This despite the Step group experiencing the full 60° rotation for only one block as opposed to three blocks for the Ramp group. The lack of disambiguation between gradual learning paradigms may have partially contributed to the conflicting findings in previous studies on gradually introduced visuomotor perturbations.

In an interesting but exploratory finding, we see not only higher implicit aftereffects of learning, but also more robust implicit aftereffects of learning in our Step group. In the Abrupt and Ramp groups, we found typical decay of implicit aftereffects over the course of multiple reaches without any visual feedback of hand position [1,7,30–34] – that is, our measures of implicit aftereffects decreased over time. In the Step group however, we find no such decay during nine No-cursor reaches (Fig 4A). Introducing perturbations in a stepped manner may cause not only larger, but also more persistent, implicit learning. Taken together, the smaller increments followed by extended consolidation may be the necessary scaffolding for increased, robust implicit learning in stepped conditions.

Given that these different gradual methods of perturbation introduction led to clear differences in implicit aftereffects of learning, we also asked whether explicit strategy use was differently affected. In line with the theory of explicit and implicit processes of adaptation being in competition with, or compensating for one another, participants in the Step group should show diminished explicit strategies compared to the ramped. In our study however, although the Step group had larger reach aftereffects during block 4 when compared to the Ramp group (21.49° vs. 11.81° respectively), we do not see any clear evidence of consequently lower explicit strategies in the Step group – or for that matter, even the Abrupt group (Fig 3C). Although a process dissociation procedure is one of many methods to measure explicit strategy use, it has been reliable in consistently detecting changes in scenarios where explicit strategy use is expected to vary [5,11,35] and it is consistent with measures of explicit strategies that ask participants to directly report aiming strategies [36]. This suggests that implicit adaptation may change in some way that is partially independent of explicit strategy use [36]. Additionally, our results challenge the commonly held assumption that adapting to gradually introduced large rotations, as in the Ramp group, is less explicit.

What features of the stepped introduction are crucial for eliciting larger, and possibly more stable implicit learning? We do not yet know. The two methods of gradually introducing a perturbation were different along two main features; the time spent ramping up to a 60° rotation, and the ability to consolidate implicit learning during adaptation. The Ramp group was introduced to a full 60° rotation in fewer trials and was not given time to consolidate learning for any given intermediate perturbation size. Although the Step group was given up to 66 trials to consolidate learning to an intermediate perturbation, is possible that shorter steps would have been sufficient to elicit more implicit learning given that saturation, and likely consolidation, of implicit components of learning can occur in far fewer than 66 trials [37,38]. In learning paradigms with stepped introductions of perturbations, along with the number of trials spent on each step, the size and number of steps may also matter. Unpublished results from our research group suggest smaller step sizes may elicit even more implicit learning than we find in our current study [39]. Additionally, measuring decay in No-cursor trials following adaptation, and perhaps across even more trials than in this study, may be useful in gauging the resilience of implicit components of adaptation. Future studies should aim to identify key features of gradually introduced perturbations that affect implicit learning. Identifying these features will allow us to maximize implicit learning, a key outcome of rehabilitation and training paradigms.

